# Attitudes, Perceptions and Barriers to Research and Publishing among Research and Teaching Staff: A Case Study

**DOI:** 10.1101/347112

**Authors:** Stanley I.R. Okoduwa, James O. Abe, Blessing I. Samuel, Aimee O. Chris, Richard A. Oladimeji, Olanipekun O. Idowu, Ugochi J. Okoduwa

## Abstract

**Background:** The economic development of any nation is centered on research. Unfortunately, research activities had suffered serious setback in tertiary research institutions in Nigeria. This study explores the attitudes, perceptions and barriers to research and publishing among academic staff in Nigerian Institute of Leather and Science Technology, (NILEST) Zaria Nigeria.

**Method:** A structured self-administered questionnaire was distributed among 130 research and teaching staff at the various Directorates in NILEST. Data are presented in frequencies and percentages for questionnaire responses.

**Result:** Exactly 93.85%of the questionnaires were validated for the study which included 81.15% male and 18.85% female. The participant were researchers (26.23%), lecturers (31.15%), Technologist (20.49%) and Instructors (22.13%). The majority of participant agreed that research is important for the institute (90.98%). A total of 81.15% believed that conducting research should be mandatory for all academic staff. Only 44.26% was self-reported to have ongoing research. Some of the obstacles reported to have prevented research activities included: lack of funding (72.13%), lack of professional mentorship (84.43%) and inadequate research facilities (89.34%). Participants without a single published paper were 54.545%. Some of the reasons given for not having published papers were “no writing experience (95.65%), high publishing fee (79.71%) and long waiting period for peer review (97.10%)”. The suggestions to improve research status by respondents included“ provision of research grants/funds (92.62%), provision of internet facilities (95.10%), mandatory publication (26.23%) and appropriate mentorship (34.43%).

**Conclusion:** Majority of the respondents believed that research is relevant, only a few were engaged in active research and published articles as evidence. It is therefore recommended that policy makers should devise strategies to focus on active research activities in order to achieve the desire research mandate and goal of institutions in the development of the nation’s economy.

## Introduction

Research is essentially the search for facts in the furtherance of knowledge. It involves the collation and analysis of information to improve human understanding of phenomena under study [1]. It entails data collection, analysis, interpretation and assessment procedures conducted in a planned manner in order to find solutions to a problem [2,3]. According to the Organisation for Economic Cooperation and Development, research is the creative work that is undertaken on a systematic basis with the purpose of increasing knowledge, and to devise new applications [4].Research is frequently carried out in tertiary and research institutions [5,6]. In most cases, research staff and graduate students are the two main groups who conduct research, the rationale being that education in tertiary institutions requires students to submit research projects, theses or dissertations for fulfillment of a degree program. Previously, lecturers were not required to conduct research on the challenges they would encounter in the course of teaching and learning [7,8]. Much reliance has been placed on experts from diverse fields like psychology, philosophy, mathematics and other sciences for the content and execution of their teaching. As a result, the contribution to the field of teaching and learning from people who were not trained as educationists has been significant and disproportionate [9].

Research was made a university function in addition to the task of teaching in the late 19^th^ century after the first academic revolution [10]. Since then, attention to research is one of the most important issues in scientific communities [1]. In recent years, research output emanating from academics, has been assessed and used to rank universities against each other [7, 11]. Publishing of research work is evidence to justify support of research investigations and a guarantee of subsequent research funding for sustainability of the institute’s mandate and organizational goals[12].However, it has been observed that the attitude and perception of both teaching and research staff of many tertiary research institutions in Nigeria is lackluster [13, 14]. Attitude and perception have significant impacts on staff performance, which in turn decides the performance of the organization. There is a need for the provision of the requirements of researchers, which would bolster their enthusiasm and improve their attitude and productivity [15].

The Nigerian Institute of Leather and Science Technology (NILEST) is a specialized agency under the Federal Ministry of Science and Technology in Nigeria with the sole mandate “To provide courses of instruction, training and research in the field of leather and leather product technology and conduct research and development on leather technologies and goods production” [16].The visions of NILEST are “to become a research institute of international standard in the provision of innovative research and development in the processing and conversion of raw hides and skins into leather and leather products. Secondly, to be a renowned center of excellence in the field of tannery effluent monitoring and control, leather and leather products technologies and lastly, to be a centre of excellence in the production of scientific models and polymer products”. The workforce of the institute comprises of both research and teaching staff employed to actualize the vision and core mandate of its establishment [16]. Unfortunately, after over 50 years of its inception coupled with the fact that the vast majority of Nigerians use leather and leather products in one way or another, the institute is virtually unknown [17].Also, there is a paucity of published literatures emanating from the institute on indexed journals, which is having a negative impact on its public profile. Possible explanations could be that research activities have been truncated for various reasons, which has a knock on effect on academic publications. It can be claimed that there is a direct relationship between conducting research and actual progress of any institute of research. Previous studies conducted in other countries mention some barriers to research to include demographic characteristics and lack of resources for publication, and these were shown to affect the attitude and perception of staff towards research work and publications [18, 19]. This study therefore focuses on the attitudes, perceptions and barriers to research and academic publishing by the research and teaching staff in NILEST, Zaria, Nigeria.

## Method

### Study Design

This study was a cross-sectional descriptive survey using a structured questionnaire. The questionnaire covered age, sex and significant attitudes and perceptions (identified through extensive literature searches as barriers to research and publishing). The questions were modified to suit the peculiar nature of the present case. The questionnaire was subdivided into different sections that included the perceptions of staff towards research, academic publishing and subsequent effects on career development. Challenges and hindrances faced by staff were also queried and collated. Research assistants were trained in basic interviewing techniques. The questionnaire was pre-tested for flow of questions and for validity and was distributed at the Nigerian Institute of Leather and Science Technology (NILEST), headquarters, Zaria, North-West, Nigeria(located on latitude: 11°9′55.3″ longitude: 7°39′5.84″).

### Ethical Consideration

The study protocol was approved by the Institutional research and academic committee of NILEST and data confidentiality, was assured.

### Selection Criteria

Employees of NILEST headquarters, Zaria, Nigeria, who are either research or teaching staff, were selected and included for the study. Non-research, non-teaching and administrative staff were excluded from the study.

### Data Source and Participants

A total of two hundred (200) staff were enrolled for the study. They were randomly selected from all the departments and units in four main directorates at NILEST headquarters, Zaria, Nigeria. The study was conducted between December 1st, 2015 and May 30th, 2016.

Based on the inclusion criteria, the final sample consisted of one hundred and thirty staff (130) from 4 directorates namely: Directorate of Leather Technology (DLT), Directorate of Polymer and Environmental Technology (DPET), Directorate of Research and Development (DRD) and Directorate of Science Laboratory Technology (DSLT). Participation was voluntary, and informed consent was obtained before distribution of questionnaires. The staff were given ample time to answer the questionnaire, after which the questionnaire were collected. Eight respondents returned incomplete questionnaire, these were considered invalid and excluded from the analysis. Only the completed questionnaires 122 (93.85%) were considered valid and coded for analysis.

### Statistical Analysis

The data obtained from the questionnaires were entered into Microsoft Excel 2013. Microsoft Excel and Statistical Package for Social Science (SPSS) software program (IBM SPSS v.20 Inc., Chicago Il, USA) were used for descriptive analysis of the data. The results are presented as percentages and frequency distribution.

## Results

### Demographic Characteristics of the Studied Participants

The percentage distribution of respondents and their demographic characteristics are presented in Table 1 and Fig 1. A total of one hundred and twenty two (122) staff participated in the study, including23females (18.85%) and 99 males (81.15%). The participants were drawn from four main directorates, as follows: 21.30% from DLT, 19.70% from DPET, 28.70% from DRD and 30.30% from DSLT. According to the designations of the participants as distributed in the four directorates, 26.23% were Researchers, 31.15% were Lecturers and 42.62% were Technologists/Instructors. The majority of the participants (42.62%) were within age range of 31-40 years. The lecturing cadre had the highest number with 31.15%. The DSLT recorded the highest number of respondents (30.30%).In terms of years of service, 64.75% of the participants were employees who had served the institute for less than 10 years. The researchers ranked highest (17.21%) among the less than 10 years of service group. It was observed that only 35.25% of the participants had served the institute above 10 years. A total of 27.05% of the participants had various postgraduate qualifications such as M.Sc./PhD, while the majority (72.95%) of the participants had only the basic first degree qualification such as B.Sc./HND (Fig1).

**Table 1:**
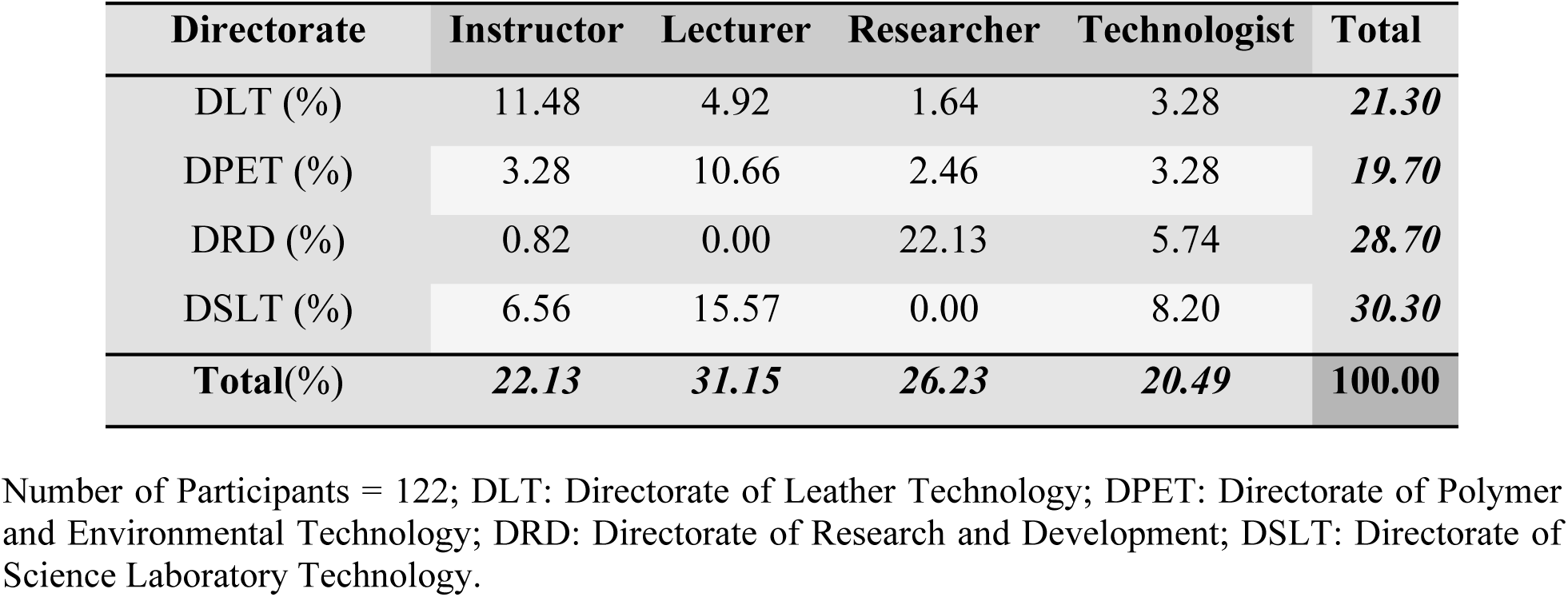
Percentage Distribution of Respondents by Designations and Directorate.

**Fig. 1:**
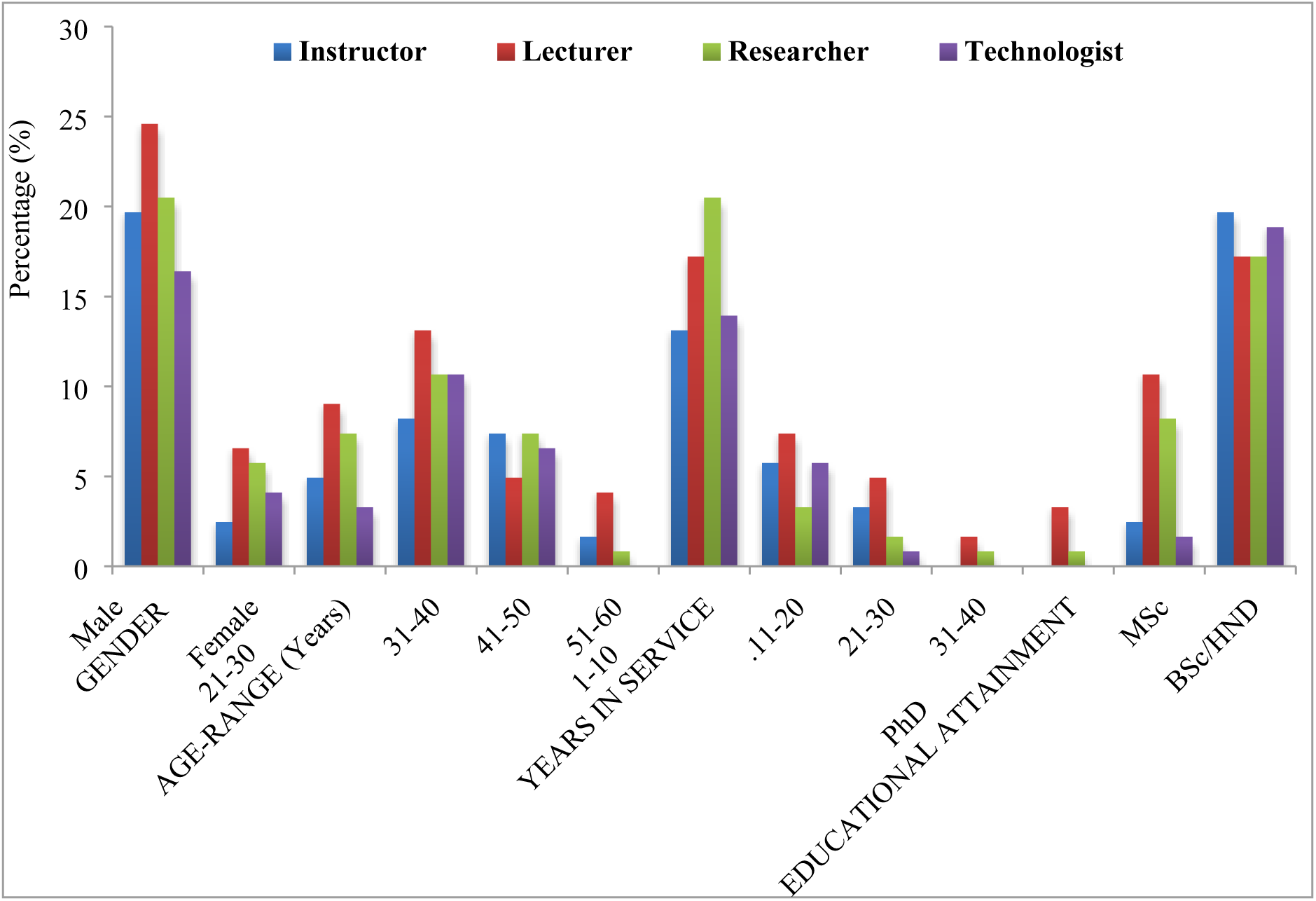
Demographic Characteristics of Respondents.

### Barriers Encountered by Participants towards Research and Publishing of Articles

Some of the specified barriers encountered by the participants in conducting research and publishing of academic articles are presented in Fig 2 and Table 2. The identified barriers included lack of research funding (72.13%), inadequate research facilities (89.34%), inadequate training/orientation programs (52.46%) and lack of mentorship (84.43%). Among the various categories of respondents, 84.38% of Researchers reported lack of funds as a barrier hindering research activities and publishing of article among them. More than 80.00% of each category of respondents reported inadequate research facilities as the most significant issue. About half of the Instructors (48.15%) reported inadequate research experience as the most challenging barrier towards research and publishing. Lack of training and orientation programmes was reported by over half of the Researchers (65.63%) and Technologists (56.00%). Lack of professional mentorship was also reported by more than 70% of each category of respondents. It was observed that majority of the respondents (79.25%) have never benefitted from research grants. Only a few of the participants (15.09%) had benefitted from publication fees assistance/waiver. None of the Instructors and Technologists has ever benefitted from publication fee assistance. Limited numbers of the Lecturers (3.77%) and Researchers (11.32%) reported that they have benefitted from publication fee assistance.

### Perceptions of Participants to Research and Publishing of Articles

The perception of respondents to research and publishing is presented in Fig 2 and Table 3. The majority of participants agreed that research and publishing is important (90.98%) for academic and economic development. From the perspective of the different groups of respondents, it was observed that all the Researchers concurred to the fact that research is important. More so, 88.89% of Instructors, 92.11% of Lecturers and 80.00% of the Technologists consented to the significance of research in increasing the research status of the institution and development of the nation’s economy. The majority of the respondents have the same opinion that conducting of research and publishing of articles should be mandatory for all staff (81.15%).Among these groups were 66.67% Instructors, 76.32% Lecturers, 96.88% Researchers and 84.00% Technologists.

**Fig. 2:**
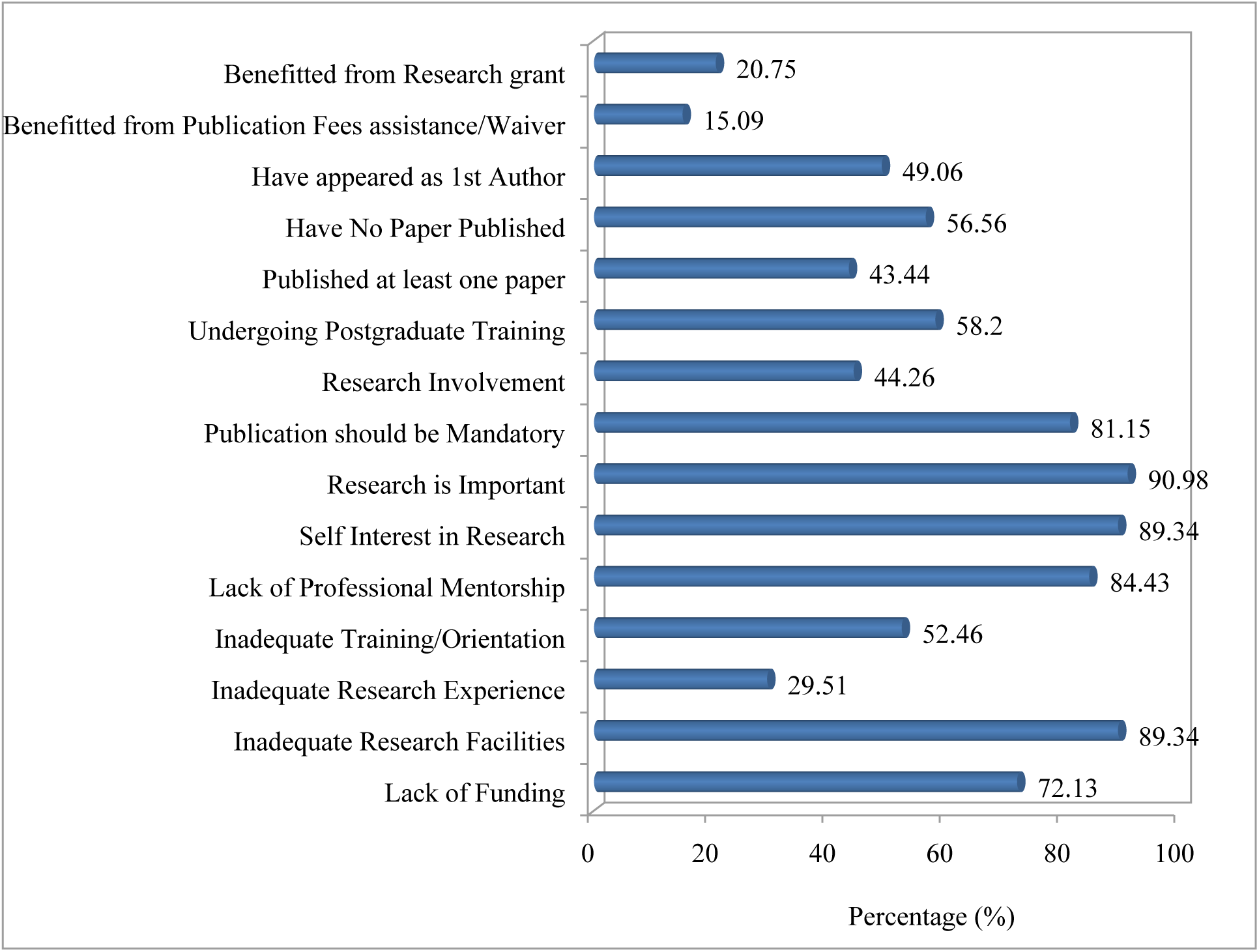
An Overview of Barriers, Attitudes and Perception of Entire Participants towards Research and Publishing of Articles.

### Attitude of Participants to Research and Publishing of Articles

The self-reported attitudes of respondents to research and publishing are presented in the Fig 2 and Table 3. Although, majority of the respondents reported that they have self-interests in research (89.34%) (Out of which 96.88% were Researchers, 92.00% were Technologists and 94.74% Lecturers and 70.37% were Instructors) it was, however, amazing to observe that only 44.26% of the participants were involved in ongoing research. Among the participants involved in active research were 75.00% of Researchers.

**Table 2:** Barriers towards Conducting Research and Publishing of Articles that are Specific among the Different Cadres of Respondents.

| Category | Respondents | Frequency | Percentage (%) |
| --- | --- | --- | --- |
| Lack of Fund<br>(72.13%) | Instructors | 20 | 74.07 |
|  | Lecturers | 29 | 76.32 |
|  | Researchers | 27 | 84.38 |
|  | Technologists | 12 | 48.00 |
| Inadequate<br>Research Facilities<br>(89.34%) | Instructors | 22 | 81.48 |
|  | Lecturers | 36 | 94.74 |
|  | Researchers | 28 | 87.50 |
|  | Technologists | 23 | 92.00 |
| Inadequate<br>Research<br>Experience<br>(29.51%) | Instructors | 13 | 48.15 |
|  | Lecturers | 10 | 26.32 |
|  | Researchers | 6 | 18.75 |
|  | Technologists | 7 | 28.00 |
| Lack of<br>Training/Orientation<br>(52.46%) | Instructors | 12 | 44.44 |
|  | Lecturers | 17 | 44.74 |
|  | Researchers | 21 | 65.63 |
|  | Technologists | 14 | 56.00 |
| Lack of Professional<br>Mentorship<br>(84.43%) | Instructors | 25 | 92.59 |
|  | Lecturers | 27 | 71.05 |
|  | Researchers | 29 | 90.63 |
|  | Technologists | 22 | 88.00 |
| Benefitted from<br>Publication fees<br>assistance/waiver<br>(15.09%) | Instructors | 00 | 0.00 |
|  | Lecturers | 02 | 3.77 |
|  | Researchers | 06 | 11.32 |
|  | Technologists | 00 | 0.00 |
| Benefitted from<br>Research Grant<br>(20.75%) | Instructors | 2 | 3.77 |
|  | Lecturers | 7 | 13.21 |
|  | Researchers | 2 | 3.77 |
|  | Technologists | 0 | 0.00 |

**Table 3:** Attitudes and Perceptions towards Conducting Research and Publishing of Articles that are Specific among the Different Cadres of Respondents.

| Category | Respondents | Frequency | Percentage (%) |
| --- | --- | --- | --- |
| Self Interest in Research (89.34%) | Instructors | 19 | 70.37 |
|  | Lecturers | 36 | 94.74 |
|  | Researchers | 31 | 96.88 |
|  | Technologists | 23 | 92.00 |
| Research is Important (90.98%) | Instructors | 24 | 88.89 |
|  | Lecturers | 35 | 92.11 |
|  | Researchers | 32 | 100.00 |
|  | Technologists | 20 | 80.00 |
| Research/Publication should be Mandatory (81.15%) | Instructors | 18 | 66.67 |
|  | Lecturers | 29 | 76.32 |
|  | Researchers | 31 | 96.88 |
|  | Technologists | 21 | 84.00 |
| Research Involvement (44.26%) | Instructors | 6 | 22.22 |
|  | Lecturers | 13 | 34.21 |
|  | Researchers | 24 | 75.00 |
|  | Technologists | 11 | 44.00 |
| Ongoing Postgraduate Training (58.20%) | Instructors | 15 | 55.56 |
|  | Lecturers | 21 | 55.26 |
|  | Researchers | 25 | 78.13 |
|  | Technologists | 10 | 40.00 |
| Published at least one paper (43.44%) | Instructors | 5 | 18.52 |
|  | Lecturers | 20 | 52.63 |
|  | Researchers | 22 | 68.75 |
|  | Technologists | 6 | 24.00 |
| Have no Paper Published (56.56%) | Instructors | 13 | 48.15 |
|  | Lecturers | 21 | 55.26 |
|  | Researchers | 25 | 78.13 |
|  | Technologists | 10 | 40.00 |
| Have appeared as 1st Author (49.06%) | Instructors | 3 | 11.11 |
|  | Lecturers | 9 | 23.68 |
|  | Researchers | 12 | 37.50 |
|  | Technologists | 2 | 8.00 |

More than half 58.20% of the Researchers, Instructors and Lecturers were undergoing various post-graduate training such as M.Sc. and Ph.D. programmes. Interestingly, only 43.44% of the participants have published at least one academic article in a peer reviewed journal. Among these were 68.75% of Researchers and 52.63% of Lecturers. Regrettably 56.56% of all the respondents have never published any paper in academic peer reviewed indexed journals. Among those that have published at least one academic paper in peer reviewed journals, only 49.06% have appeared as first author in those articles of which 37.50% were the Researchers.

Those who reported they had not published articles were questioned further as to why they had not. Some of the respondents 15.94% said it was due the long waiting period for peer review process. About 20.29% reported they had no time to write article for publication consideration due to other commitments while 33.33% said it was due to lack of departmental motivation to conduct research and consider publishing the outcome of the research. A large number 97.10% said due to rejection of manuscript upon submission for publication consideration, 95.65% said they had no writing experience and 79.71% said it was due to the high publication fee for publishing articles (Fig 3).

About half 53.62 of the respondents in the group said they had no mentorship on how to process manuscript for peer review indexed journals. Surprisingly, about 72.46% of those without published articles said it was not needed for their promotion/career growth (Fig 3).

**Fig. 3:**
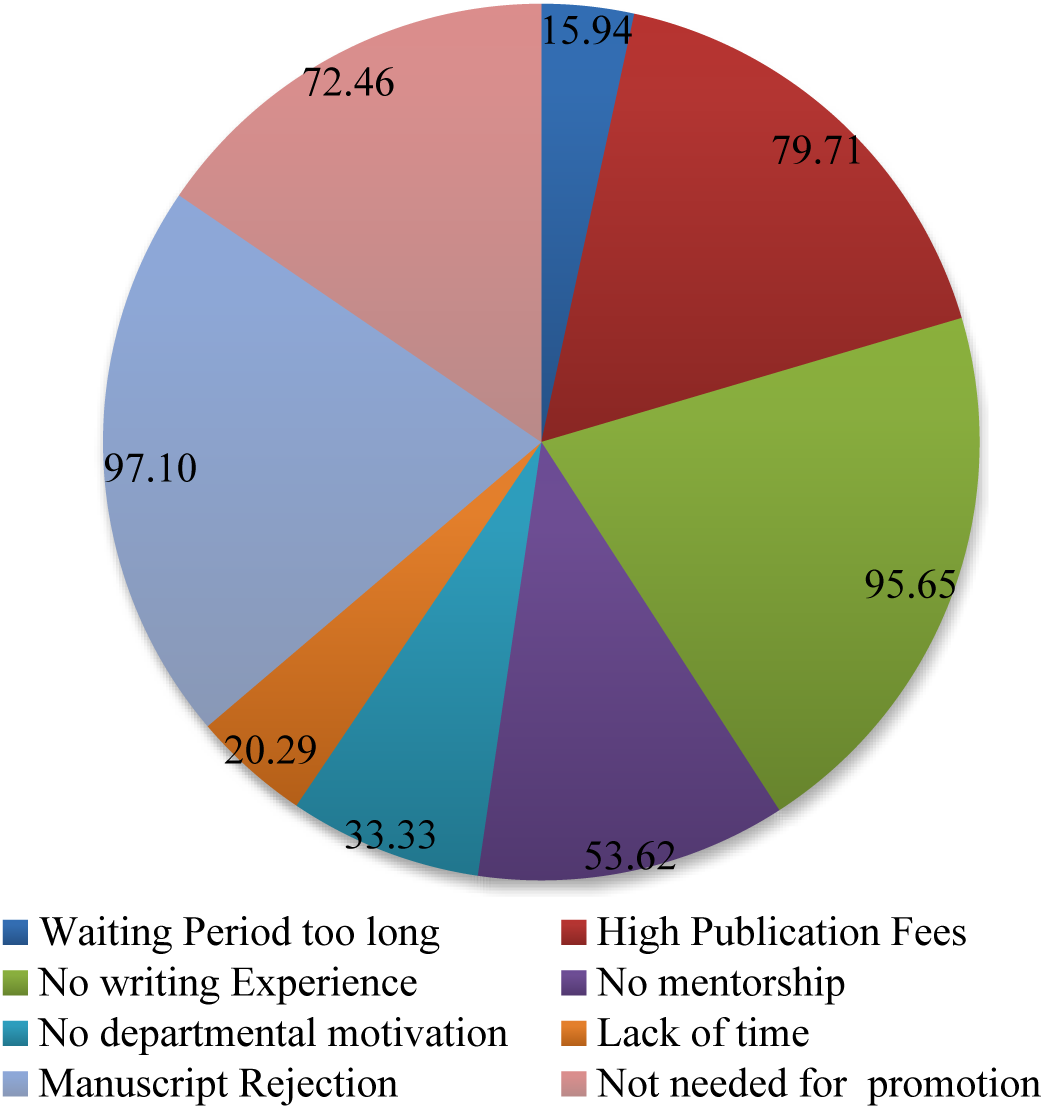
Reasons for Not Publishing by Participants without Published Article. NB: Values presented are in percentages

Those who have published at least one paper or submitted manuscript for publication consideration reported their main motivation to be career development, information relay and self-interest (Fig 4).

**Fig. 4:**
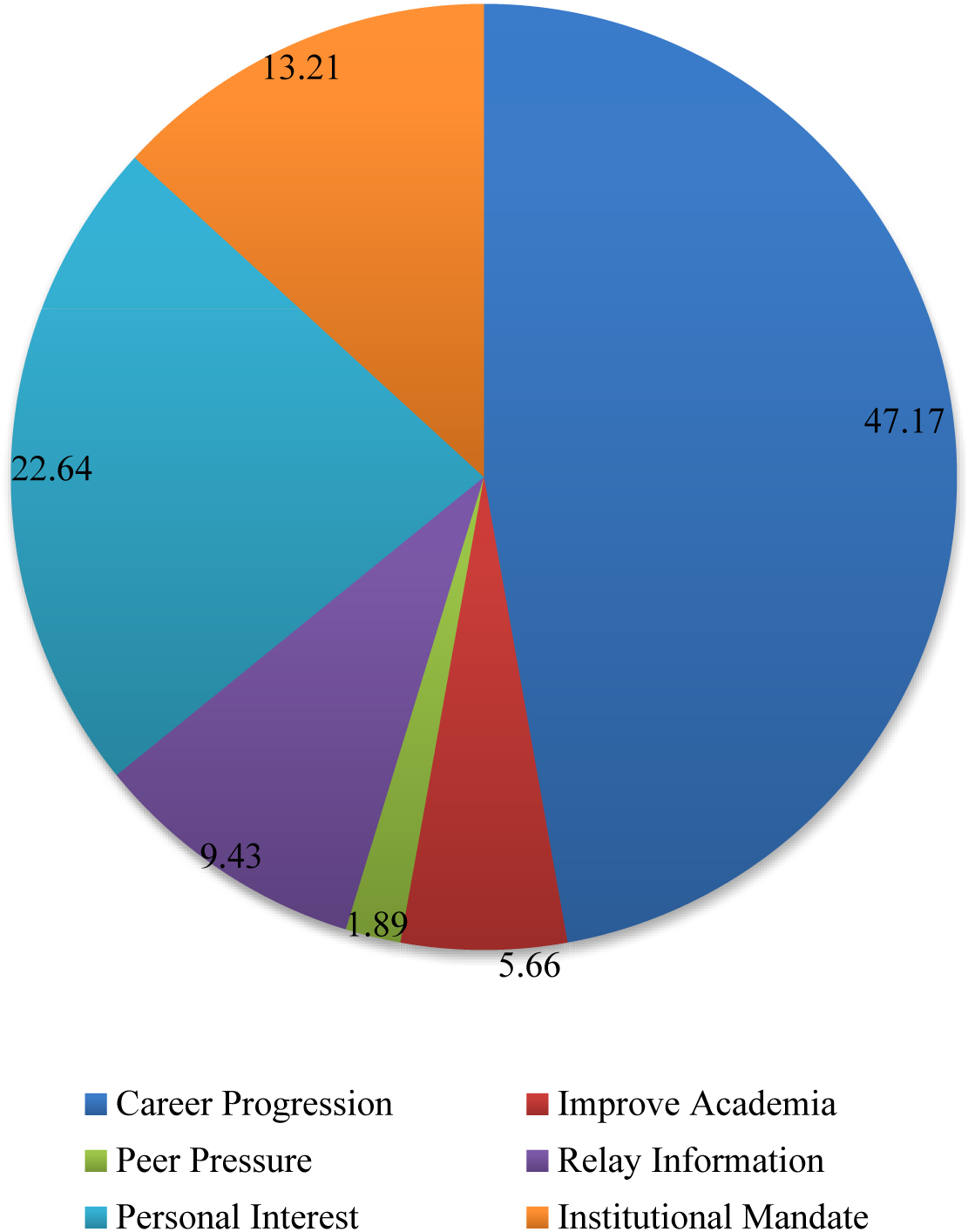
Reasons for Publishing by Participants with at Least One Published Article. NB: Values presented are in percentages

### Suggestions to Improve the Status of Research and Publishing in Tertiary Institutions

Suggestions collated from the participants that could help improve the status of research and publishing are presented in Fig 5. These included provision of research grants and publication fee assistance, special award of excellence for outstanding contributions to deserving staff and provision of internet facilities.

**Fig. 5:**
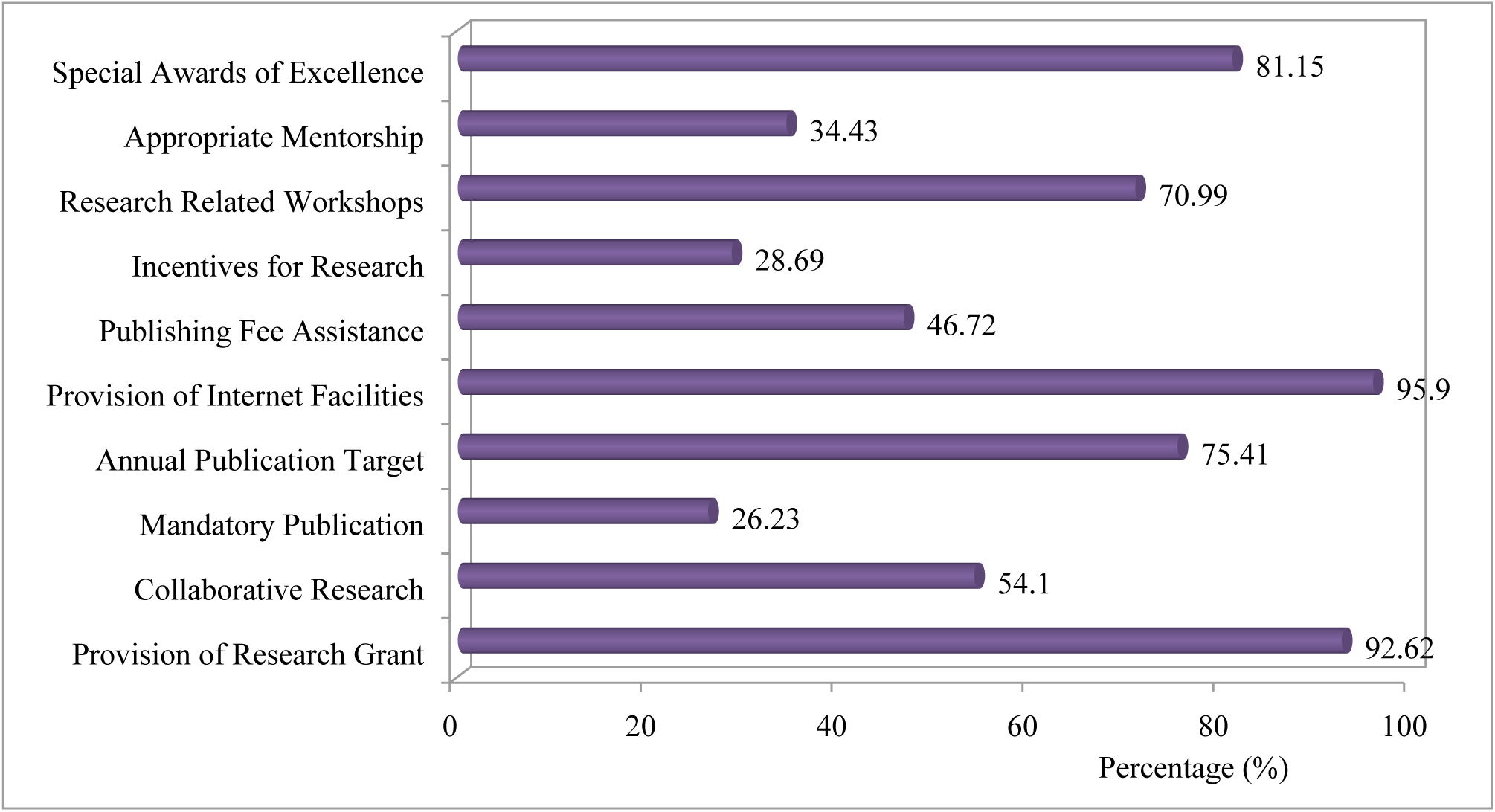
Suggestions by Participants for Improving the Status of Research and Publishing.

## Discussion

This study revealed a great disparity between the participation in and attitudes toward research activities and publication of scholarly articles among research and teaching staff in NILEST, as well as significant barriers impeding these activities. Some of which have been reported by other investigators [7,12,20,21] such as lack of adequate facilities, skills and personal interest. To the best of our knowledge this is the first study of its kind to examine the barriers, perceptions and attitudes of staff towards research and/or academic publishing in NILEST. The overall response of staff was encouraging, the initial reluctance of a sizable proportion of the staff notwithstanding. The perceptions and attitudes of staff have significant impact on the success of an organization [22, 23]. More so, staff satisfaction towards research and publishing practices will enhance the developmental goal and strategic management of the institution and vice versa [24, 25]. The attitude and perception of staff towards research and publishing in the institution essentially depends on job satisfaction [26]. Job satisfaction of staff in turn is influenced by research and publishing practices; research funding, research leave, research allowances, research training and development compensation, availability of amenities and professional mentorship [23,25].

Although, gender equality is an imperative consideration globally [27, 28] this study showed that the number of male working as research/teaching staff was higher compared to females in the entire directorate investigated (Fig 1). Among the female staff, involvement in research was notably lower compared to males. This observation is similar to some studies conducted in the US, one of which cites low self-ability as a major barrier toward participation in research activities. However, no discrete reason was recognized in our study to elucidate this observation. On the other hand, this difference may be due to the cultural, religious and social expectations and responsibilities faced by females in this part of the world [29, 30].

The majority of respondents in our study had been in active service with the institute for less than 10 years. This may be due to a change in the name of the institute that properly reposition it in line with its mandate, mission and vision, which occurred on 1^st^ April 2011 (7 years ago) [16,31]. This change brought an influx of research oriented staff, in line with the new research capable status of the institute (it was previously a college).It was earlier reported that there were just 2 Ph.D. and few numbers of M.Sc. holders as of 2009. But after the change in name from CHELTECH to NILEST the numbers increased to 7 Ph.D. and 15 M.Sc. holders excluding additional 5 Ph.D. and 7 M.Sc. staff in training as of 2012 [16].

A probable reason for the observed decline in staff strength as the years of service increases may be a lack of job satisfaction, leading to transfers of service to perceived greener pastures. This is most likely responsible for the dearth of Ph.D. holders in the institute, as upon attainment of higher qualifications, there is the urge to transfer service to universities where staff may be better placed and accorded portfolios befitting their status. This may include professorial seats and administrative or political positions. This observation may be supported by the fact that more than half (58.2%) of the participants were engaged in various postgraduate programmes, yet only a few (less than 5) hold a doctorate degree as of the time of this investigation. In the Nigerian system it takes around10 years for a civil/public servant to acquire a doctorate [2 year confirmation, 2 year study leave (MSc), 2 year waiting period, 3 years study leave (Ph.D.)]. The greener pastures beckon thereafter.

Some other barriers to research activities identified by this study have been described by other investigators elsewhere [32-34]. These include lack of suitably qualified mentors with appropriate expertise and sufficient time for mentoring, limited resources such as funds and facilities and logistical difficulties. In all, lack of support from the institution and inappropriate funding of research activities are the hallmarks of failure to execute research tasks. Besides these, the limited number of research mentors is one of the barriers claimed by some of the respondents in this study. Lack of mentorship contributes to the inadequate research experience confronting the staff [35-38]. Lack of professional mentorship advanced as an incredibly strong theme predominantly for new members of academic staff in this study who acknowledged the need for guidance in commencing research works. Positive role models and adequate professional mentorship are crucial to researchers and, if unsupported, early career staff can discontinue their work [34]. It would be rational to believe that the best mentor would be an experienced researcher with track record of publications. This observation was in harmony with the report of Williams [7] on the need for new researchers to have a mentor who can guide them through the process of research and publishing of scholarly articles. Unfortunately, this practice is only obtainable in tertiary institutions where research is part of students’ academic program of study [33].

The majority of the respondents (90.98%) agreed that it was imperative to publish papers. This observation is consonant with reports from previous investigators on related subjects [12,39]. All the Researchers included in this study felt that research activities and publishing are mandatory for career progression, a condition of service for research institutions in Nigeria. This is contrary to the conditions of service in use by the teaching staff in NILEST that provides no recognition for research activities and academic publications. The respondents’ interest towards research-and reasons for which scholarly publication was considered valuable-included improving their relationships with and gaining respect from fellow colleagues in the scientific community, advancing their career opportunities, and improving their writing and research skills. These observations were similar to that reported by Griffin and Hindocha [40]. Research skills for teaching staff are becoming imperative, particularly for obtaining designated positions in competitive markets, and in order to secure research grants [41]. Aside research staff, teaching staff can be potential contributors to scientific research and development through participation in different commercially oriented research [42,43]. It is noted in this study that less than half of the respondents (43.44%) reported scholarly research publications as a means to improve the relationship with and gain respect from scientific community, as well as recuperating their writing and research skills.

For those that had not published at least one article, it is clear that the main barrier was not having the opportunity to perform research as they were not engaged in any research activities in the first instance; hence, they feel they have nothing to declare as a publication. This was also in agreement with the findings reported by Griffin and Hindocha [40]. A survey of Australian researchers showed that research infrastructure support is vital to research productivity[44].

In this study, 81.15% of the respondents have suggested that research and publishing should be made mandatory for all academic and research staff (Table 3). Mandatory involvement in research activities has been demonstrated to improve researchers’ attitude towards research [45].

Lack of time is another factor militating against conducting of research and also one major hindrance to accessing research funds [32]. According to Yusuf, [46] the declining research productivity in the Nigerian university system is attributed to among other factors the “rising workloads associated with deteriorating staff/students ratio, which leave little time for research. Lack of knowledge about funding agencies/organization, bureaucracy in the acquisition of funds and the exclusion of research institutes from Education TrustFund by the 2011 *Tertiary Education Trust Fund (TETFund) Acts*, in which section 20 of the Act deterred researchers from conducting meaningful research[32,47]. These observations were in agreement with the findings by Dorsey *et al*., [48] that government policies on budgets to researchers deter implementation and conducting of research.

Due to the anonymous nature of the present research, the likelihood of bias is reduced. This does not sideline the significant limitation, which need to be highlighted. The present study was conducted in a single research institute, hence may be relative when one has to generalize.

## Conclusion and Recommendation

In conclusion, this study highlights clearly that less than half of the studied population were involved in active research activities and had published at least one article. This can be improved by providing more accessible opportunities to take part in research followed by encouragement and guidance from professional mentors. More opportunities to perform research and teaching will aid academic staff to increase their publishing potential, Hence, the management of the institute and policy makers may consider taking special research initiatives to address the barriers and improve the involvement of academic staff in scholarly active research activities and publishing.

The results of this study could be the basis for similar future comparative studies and policy formulations that may promote the research mandate of every institution in order to achieve their organizational goal in enhancing the nation’s economic development, security and sustainability. Further study is therefore recommended at different research institutions on a larger sample size.

## Acknowledgments

This research did not receive any specific grant from funding agencies in the public, commercial, or not-for-profit sectors. The authors express their warm appreciation to the Dr. A. Ejila, Director of Research, NILEST, Zaria, Nigeria for approval to conduct the study.

